# Anti-HIV Immunotoxin and Antibody-Drug Conjugate Display Both Common and Distinct Effects in Killing Target Cells

**DOI:** 10.64898/2026.04.07.717054

**Authors:** Seth H. Pincus, Tami Peters, Megan Stackhouse, Galen O’Shea-Stone, Frances M. Cole, Brian Tripet, Valérie Copié

## Abstract

**Background:** In the course of testing mAb-based therapies to eradicate the persistent reservoir of HIV infection, we investigated the efficacy and mode of killing of HIV-infected cells by two categories of cytotoxic immunoconjugates (CICs) targeted by the same mAb, an immunotoxin (IT) and antibody-drug conjugate (ADC).

**Methods:** We performed metabolic and transcriptional analyses of treatment effects on the persistently-infected cell line H9/NL4-3. Cells were treated with CIC’s consisting of the anti-gp41 mAb 7B2 conjugated to either deglycosylated ricin A chain (dgA) or to the highly cytotoxic anthracycline derivative PNU-159682. At intervals up to 24 hr, intracellular metabolites were quantified by ^1^H nuclear magnetic resonance spectroscopy, and the transcriptome analyzed by RNA-Seq.

**Results:** Six hr post treatment, 7B2-dgA elicited both metabolic and transcriptional alterations, whereas 7B2-PNU treated cells did not differ from untreated cells. 7B2-dgA treated cells exhibited elevated intracellular levels of many amino acids, and activation of gene pathways for apoptosis, intracellular signaling, and immune activation. By 24 hr, both 7B2-dgA and 7B2-PNU treated cells differed markedly from untreated. Many of the changes observed following 7B2-PNU treatment at 24 hr were similar to those observed at 6hr following 7B2-dgA, likely indicating processes involved in cell death, but a number of alterations were unique to either IT or ADC treated cells.

**Conclusions:** An IT and ADC showed both similarities and differences in their cytotoxic effects. These results raise the question of whether the mode of cell killing could be a determinant of clinical efficacy. Although these studies were aimed at targeting the persistent reservoir of HIV infection, they have relevance for the design of CICs to treat cancer and other conditions.

**SUMMARY:** The use of cytotoxic immunoconjugates, wherein an antibody is attached to a cellular poison, is effective in the treatment of cancer and other conditions. We seek to extend these results to treating HIV and other chronic viral infections. We analyzed the molecular mechanisms of cell killing when the same antibody was attached to different toxic structures. We report that each immunoconjugate induced both common and distinct patterns of killing. Such differences may have clinical relevance.

## INTRODUCTION

To eradicate the reservoir of HIV infection that persists in patients despite effective antiretroviral therapy (ART), the construction of cytotoxic immunoconjugates (CICs) to eliminate HIV-infected cells has been pursued by our group (1–10) and others (11–15). Our initial studies were performed with immunotoxins (ITs), which were effective both *in vitro* and *in vivo,* but proved highly immunogenic (4–10). Using the ITs, we identified the helix-loop-helix region of gp41 as the most effective target of CICs requiring internalization to be effective (3, 4, 7, 9). Following the lead set in oncology, we then produced antibody drug conjugates (ADCs) as an attempt to mitigate immunogenicity. Our initial constructs utilized small drugs found effective in oncology, but yielded anti-HIV ADCs lacking *in vitro* efficacy (5). We subsequently identified PNU-159682, an anthracycline derivative 6000X more potent than doxorubicin, as producing an ADC that is cytotoxic to HIV-infected cells and has antiviral effects (1).

Because of the immunogenicity of ITs, the clinical uses of ADCs in oncology far outstrip those of ITs. Other than immunogenicity, it has not been shown whether the use of ITs vs ADCs differ in other ways that may affect their clinical utility. To address this question we have constructed an IT, an ADC and a radioimmunoconjugate (RIC) targeted by the same mAb to HIV gp41 and compared their cytotoxic and antiviral efficacy in the same experimental systems (1, 16, 17). The IT was 7B2-dgA (deglycosylated ricin A chain), a well characterized IT with efficacy in macaques (5); the ADC was 7B2-PNU (1); and the RIC was 7B2-^225^Ac, a high energy alpha emitting isotope. We have found that the IT was more potent than the ADC (IC_50_ on Env+ 92UG transfectant cells: 7B2-dgA 0.56 ng/mL, 7B2-PNU 12.4 ng/mL), with both having equivalent non-specific toxicity on the Env-negative 293T parental cells (IC_50_ 3100 vs 3300 ng/mL). Killing was more rapid with the IT and RIC than the ADC. In cultures containing mixtures of Env+ and Env-negative cells, ADC and RIC resulted in bystander killing of Env-negative cells, the IT did not. Low doses or short exposure to the ADC resulted in enhanced cell division (17).

In this manuscript, we further characterize the differences between 7B2-dgA and 7B2-PNU by examining the effects of the CICs on HIV-infected cells at a molecular level, performing a metabolic and transcriptional analysis over the initial 24 hr following cytotoxic therapy. These interrelated parameters were chosen because they reflect the dynamic processes the cells were undergoing following cytotoxic treatment. The analyses again point to differences in kinetics of the response. In the process of dying, the cells demonstrated common responses, including activation of apoptosis and canonical cell-signaling and immune activation pathways, as well as CIC-specific pathways. The clinical relevance of the observed differences between IT and the ADC remain to be demonstrated. There has been considerable effort expended in “de-immunizing” ITs with moderate efficacy (5, 18–24). Should these efforts attain success, our results suggest that ITs could yet achieve a place in the therapeutic arsenal.

## MATERIALS AND METHODS

### Reagents and cell lines

MAb 7B2 binds CSGKLIC, the conserved 7 amino acid loop within the helix-loop-helix region of HIV gp41 (25). 7B2 was originally produced by James Robinson (Tulane University) and purified 7B2 was the gift of Barton Haynes (Duke University). 7B2-dgA consisted of deglycosylated ricin A chain (dgA), the kind gift of Ellen Vitetta (University of Texas Southwestern), conjugated to 7B2 using the heterobifunctional cross-linking agent succinimidyl 6-[3(2-pyridyldithio) propionamido] hexanoate (SPDP, Pierce Biotechnology, Rockford, IL USA) at a 1:1 mass ratio as described previously (5). Deglycosylating ricin A chain avoids non-specific toxicity of A chain-associated glycan that is bound by mannan receptors on hepatocytes (26). Products were characterized biochemically by SDS-PAGE and contained a mixture of 0, 1, 2, and 3 dgA per 7B2. 7B2-PNU was produced at Levena Biopharma as described elsewhere (1). The preparation contained a mixture of 0, 1, 2, and 3 PNUs per 7B2, similar to that of 7B2-dgA. It should be noted that the linker in 7B2-dgA contains a disulfide bond which may be cleaved, whereas the linker in 7B2-PNU is non-cleavable. CD4-IgG2 was a gift from Progenics Laboratories and Paul Maddon. It consists of a human IgG2 with the V-regions replaced by the N-terminal domain of CD4 (27). sCD4_183_ is a monomer consisting of the N-terminal domain of human CD4 and was a gift from Upjohn Laboratories.

H9/NL4-3 cells are H9 CD4+ lymphoma cells persistently infected with the NL4-3 molecular clone of HIV-1 (28). These non-adherent cells maintain a productive infection in virtually all cells over prolonged passage (3, 4, 9), likely resulting from a truncation of the Vpr gene in the single integrated provirus (GenBank: AF070521.1). H9/NL4-3 cells were the gift of Kathy Wehrley (NIAID Rocky Mountain Laboratories). Cells were cultured at 37° in a humidified CO_2_ atmosphere and maintained by serial passage not to exceed a concentration of 10^6^ cell/mL in RPMI 1640 medium (GIBCO, Thermo Fisher, Waltham MA USA) with 10% fetal bovine serum (Hyclone, Logan UT USA), and penicillin/streptomycin (GIBCO).

### Cytotoxicity assays

Cell viability was assayed by trypan blue dye exclusion, which requires membrane integrity, and by MTS dye reduction, a measure of oxidative phosphorylation. Trypan blue dye exclusion, in 0.2% trypan blue (Sigma Aldrich, St. Louis, MO, USA) is semi-quantitative. MTS dye reduction was performed in 96 well flat-bottom plates in triplicate. In each well, 2 X 10^4^ H9/NL4-3 cells were suspended in 0.2 mL RPMI 1640 + 10% fetal bovine serum (FBS) in the presence of the CIC at the indicated concentration and 500 ng/mL of CD4-IgG2 (27). We have shown that the presence of soluble CD4 markedly enhances the efficacy of anti-gp41 CICs by increasing both expression and internalization of cell-surface Env (2, 8). Cells were incubated for 72 hr. During the final 3 hr of incubation, MTS/PMS substrate (CellTiter AQueous, Promega, Madison WI USA) was added. Absorbance (A) at 490 nm was read on a microplate reader (BioTek, Winooski, VT USA). The percent cytotoxicity was calculated according to the formula below:

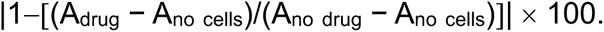

### Immunofluorescence to quantify Env expression and Annexin V binding to H9/NL4-3 cells

To measure cell-surface gp41, H9/NL4-3 cells (2 X 10^5^) were incubated with 10 µg/ mL of anti-HIV-1 gp41 mAb 7B2 + 500 ng/mL of sCD4_183_ in PBS, plus 1% bovine serum albumin and 0.1% sodium azide for 45 min at room temperature. Cells were washed twice in PBS and incubated with 2 µg /mL of secondary FITC-conjugated goat anti-human IgG (Novus Biologicals, Centennial CO USA) for 1 hr at room temperature in the dark, then washed a final time with PBS and resuspended in 2% paraformaldehyde. Apoptosis was detected in unfixed cells by binding to Annexin V-FITC (Apoptosis Detection Kit, eBioscience, San Diego CA USA) following the manufacturer’s instructions. H9/NL4-3 cells were incubated with 7B2-dgA (125 ng/mL) plus CD4-IgG2 (500 ng/mL), 7B2-PNU (500 ng/mL) plus CD4-IgG2, or no agent under standard tissue culture conditions. Aliquots were removed at 6, 14 and 22 hr and stained for binding to annexin V. FITC-stained cells were analyzed using a BD C6 flow cytometer (BD Biosciences, Franklin Lakes NJ USA) and FloJo software (BD Biosciences).

### Metabolic analyses by ^1^H nuclear magnetic resonance spectroscopy

Persistently infected H9/NL4-3 cells were grown in RPMI 1640 medium + 10% FBS without a CIC, or with 7B2-dgA 62.5 ng/mL + CD4-IgG2 500 ng/mL, or 7B2-PNU 250 ng/mL + CD4-IgG2 in 12-well culture plates (4mL per culture) for the indicated periods of time. Initial cell densities were adjusted to yield equivalent numbers of cells at 5-8 X 10^5^ cells/mL at the time of harvest, based on the observed 18 hr doubling time. Cell count and viability were determined when the cells were harvested. Cells were then washed 2X in PBS and the cell pellet frozen. Polar metabolites were separated and extracted from cell lysates using a methanol/chloroform/water solvent mixture. Metabolites were identified and quantified by proton nuclear magnetic resonance (^1^H NMR) spectroscopy, as we have described in detail elsewhere (29). The NMR spectra were processed with Bruker TOPSPIN software, followed by use of the Chenomx Software Suite for additional spectral processing including baseline correction and spectral phasing, and for identification and quantification of polar metabolites. Sodium trimethylsilylpropanesulfonate (DSS) was used as an internal standard for metabolite quantification and chemical shift referencing, while imidazole was used to correct for signal shifts arising from pH variations among samples. Tables of metabolite concentrations were generated from analysis of ^1^H NMR spectra using Chenomx and normalized to mean, resulting in final concentration tables reported in µM (Supplementary Table S1). The metabolite concentration tables were uploaded to the MetaboAnalyst v4.0 web server where concentrations of metabolites were log-transformed and auto-scaled (based on mean divided by the standard deviation of the variables) prior to univariate and multivariate statistical analysis. Student t-test, analysis of variance (ANOVA) and principal component analysis (PCA) were performed to identify potentially distinct treatment-associated metabolite patterns using Metaboanalyst v4.0, as well as in house an in-house Python workflow (pandas, statsmodels, SciPy, statannotations, matplotlib; available at https://github.com/Galenosheastone/Copie_Group_Streamlit_Stats_Tools). One-way ANOVA applying Tukey’s post-hoc analysis was performed to determine which metabolites showed statistically significant (p<0.05) differences in concentration based on treatment of cells.

### Transcriptional analyses by mRNA sequencing

H9/NL4-3 cells were grown in the presence of 7B2-dgA or 7B2-PNU with CD4-IgG2, or their absence, as described for the metabolic studies. At 6 and 24 hr cells were removed by aspiration, spun, washed once in PBS and the cell pellet frozen. Total RNA was prepared from frozen cell pellets using RNeasy plus Mini Kits (Qiagen, Germantown MD USA), following the manufacturer’s instructions; a small aliquot was removed to measure concentration (NanoDrop, Thermo Fisher Scientific, Waltham MA USA) and assess quality (TapeStation, Agilent Technologies, Santa Clara, CA), and the remainder of the sample frozen at -80° until shipment. The construction and sequencing of cDNA libraries from total RNA was performed at Azenta Life Sciences (South Plainfield NJ USA). Sequencing was performed on Illumina HiSeq sequencers with bidirectional reads. Raw data was provided to us in the form of fastq, bam, and bed files. Processed data were provided to us in the form of hit tables (only unique reads that fell within exons were included), quality control reports, and comparative analyses performed by DESeq2 (Bioconductor.org). All samples exceeded Azenta’s QC thresholds for read quality. In analyzing the data appears in this publication, we utilized the hit tables normalized to transcripts per million (TPM) and filtered out transcripts with <10 copies or a variance >15%. Data was analyzed with Microsoft Excel, MacVector (Version 17, MacVector Inc, Apex, NC USA) and ExpressAnalyst (30). The latter incorporates a variety of publicly available packages, including DESeq2 and GSEA. Normalized hit tables were uploaded to ExpressAnalyst, log transformed and auto-scaled prior to analyses. Gene pathway analysis (GSEA) was performed using tools provided by the Kyoto Encyclopedia of Genes and Genomes (KEGG, genome.jp/kegg/pathway), Reactome pathway database (reactome.org) and the Gene Ontology Resource (geneontology.org). Adjusted p-values (padj) include a strict Bonferroni correction for multiple test comparisons. Transcriptomic datasets have been deposited in the Gene Expression Omnibus (GEO, National Center for Biotechnology Information), entry number GSE289396.

### Data display and statistics

Unless otherwise stated, all samples were prepared and analyzed in triplicate. Results are shown as mean and SEM. If no error bars are visible, then the symbol on the graph is larger than the error. We have color coded the immunoconjugates as follows: blue for negative control (either unconjugated 7B2 or no treatment), gold for 7B2-dgA, and red for 7B2-PNU. Calculations were performed and graphs drawn using Microsoft Excel and GraphPad Prism v9 (GraphPad, Boston MA USA). Statistical analyses of the metabolomic and RNAseq data were as described above.

## RESULTS

### Characterization of effects of CICs on HIV-infected H9/NL4-3

H9/NL4-3 T-cell lymphoma cells are persistently infected with the HIV isolate NL4-3 (4, 28, 31). Greater than 95% of cells constitutively produce infectious HIV. Figure 1 demonstrates the interaction of H9/NL4-3 with anti-gp41 mAb 7B2 and its derivative CICs. Panel A establishes expression of the 7B2 target epitope on the cell surface, with ∼100-fold increase in fluorescence compared to control. Panel B displays a dose-response of cytotoxicity of the CICs + CD4-IgG2, with 7B2-dgA demonstrating an IC_50_ of 8.9 ng/mL and 7B2-PNU at 105.6 ng/mL, indicating that the IT is considerably more potent than the ADC on infected H9/NL4-3 cells, as it was on transfected 92UG cells (1). We have reported that infected H9/NL4-3 cells were less sensitive to the effects of the CICs than 92UG transfectants (1–3), and we observed that here as well. Panel C indicates that apoptosis, as measured by annexin V staining, was slower to develop following treatment with 7B2-PNU than with 7B2-dgA, again supporting previous observations of more rapid killing by the immunotoxin (1, 17). We have previously demonstrated the specificity of CIC killing of H9/NL4-3 cells by showing the absence of effects of CIC on uninfected H9 cells, of unconjugated 7B2 or sCD4 on H9/NL4-3, or irrelevant CIC on H9/NL4-3 (1–5, 9, 10, 32).

**Figure 1.**
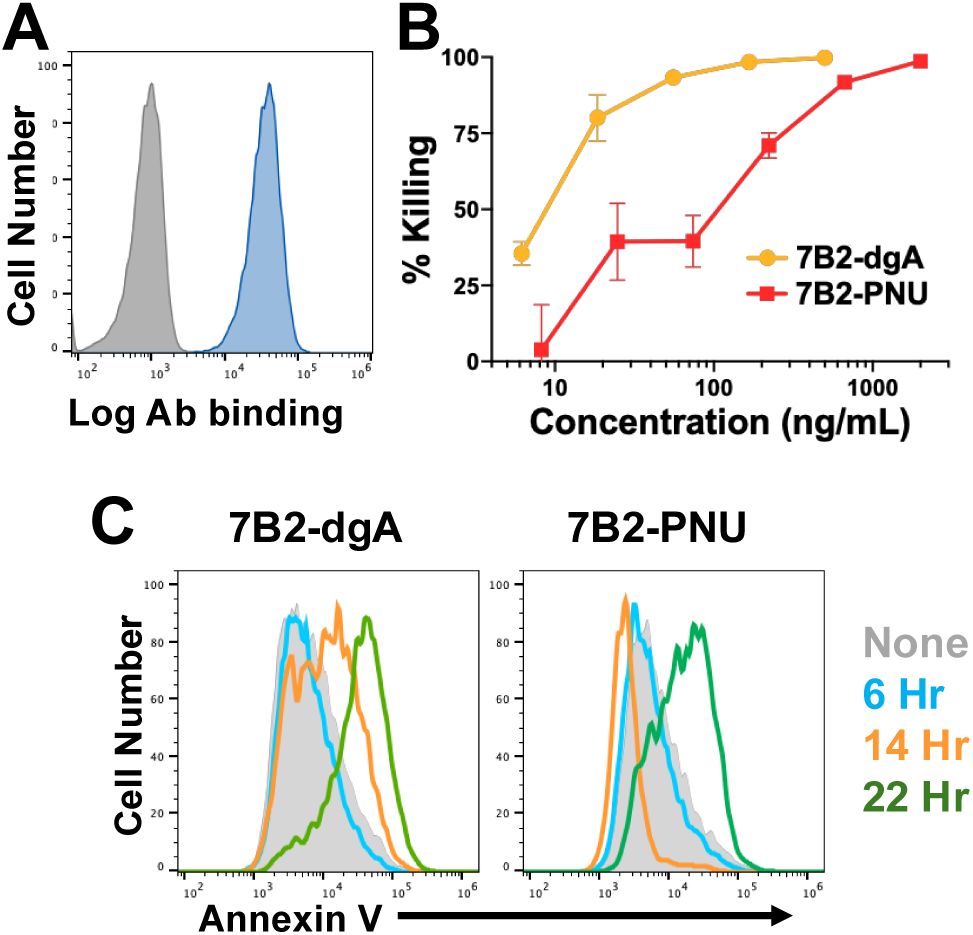
Epitope exposure and CIC-mediated killing of H9/NL4-3 cells. **A**. The binding of anti-gp41 7B2 to H9/N:L4-3 cells was demonstrated by indirect immunofluorescence and flow cytometry. The gray histogram indicates binding in the presence of secondary FITC-conjugated goat anti-human IgG only, the blue histogram demonstrates fluorescence of cells first incubated with 7B2 followed by FITC-conjugated secondary antibody. **B.** Comparative cytotoxicity of 7B2-dgA and 7B2-PNU. Cells were incubated with the indicated concentration of CIC and CD4-IgG2 (500 ng/mL) for 72 hr. MTS dye reduction was measured during the final 3 hr of culture. **C.** The kinetics of apoptosis induction by the CICs was measured by the binding of fluorescent annexin V to cells at different times following treatment with CIC.

### Metabolic effects of CIC treatment

To determine the effects of treatment with the different CICs on metabolites within the H9/NL4-3 cells, we employed ^1^H NMR spectroscopy to quantify polar metabolites (29) at selected times following either no treatment, or exposure to either 7B2-dgA or 7B2-PNU and CD4-IgG2. At each time point, prior to metabolite extraction, viable cell counts were performed using trypan blue. Growth arrest was apparent in the 7B2-dgA treated cells at 12 hr and in 7B2-PNU at 24 hr. At 24 hr, the percentage of viable cells had begun to drop, more so after 7B2-dgA than 7B2-PNU treatment. After two washes with cold PBS, intracellular metabolites were extracted and the resulting metabolite patterns analyzed.

Thirty-one metabolites were identified and quantified. Metabolite concentrations are reported in Supplementary Table S1. Figure 2A represents a principal component analysis (PCA) performed on the data. At 6 hr, 7B2-dgA treated cells displayed markedly different metabolite profiles than untreated controls and 7B2-PNU treated cells, with the latter two groups completely overlapping in the 2D PCA scores plot. At 12 and 24 hr, the metabolite patterns of 7B2-PNU treated cells gradually separated from untreated cells, while 7B2-dgA treated cells remained quite distinct. These data again highlight the difference in the kinetics of cellular responses to the different CIC treatments. Some metabolite changes in treated cells at 12 and 24 hr may have resulted from leakage of intracellular metabolites through the porous membranes of dying cells. Figure 2B indicates that the mean concentrations of almost all intracellular metabolites drop in 7B2-dgA treated cells with time, but not in untreated or 7B2-PNU treated cells. In the 7B2-dgA group, total drops in metabolite concentrations of 20% between 6 and 12 hr, and 33% between 6 and 24 hr were observed. For this reason, metabolite profiles of 7B2-dgA treated cells were only analyzed at 6 hr. It should be noted that in all treatment groups there were marked time-dependent increases in glucose and lactate concentrations and decreases in acetate and butyrate levels.

**Figure 2.**
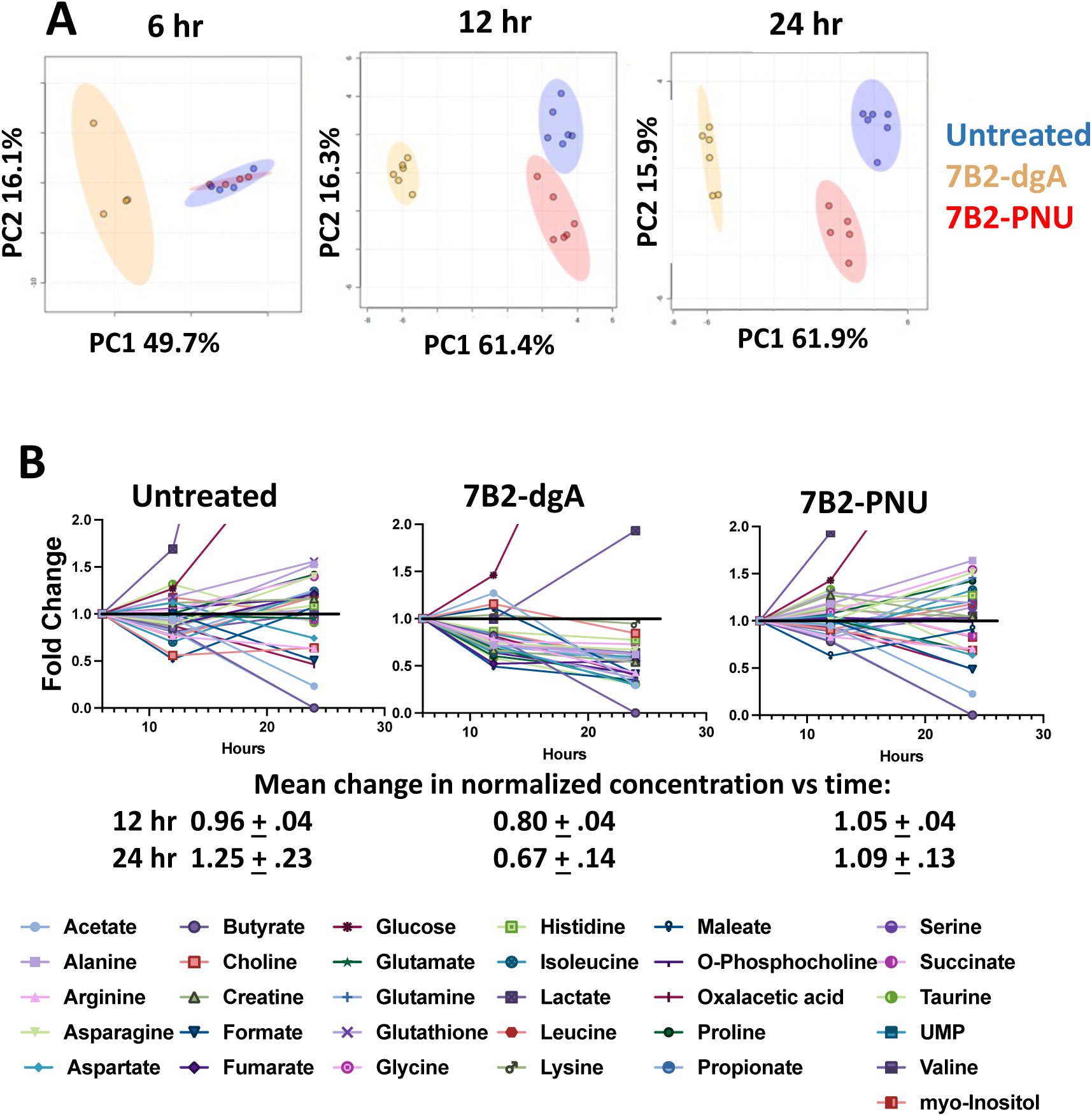
Analyses of metabolite concentrations at different times post-treatment with CIC. Replicate cultures were incubated with CIC and harvested at the indicated time points. Intracellular metabolite concentrations were determined. **A.** 2D PCA scores plots were performed using MetaboAnalyst 4.0. Data normalization and scaling parameters included: normalization to constant sum, data transformation to Log10; autoscaling (mean-centered, divided by standard deviation). **B.** Mean concentrations of each metabolite were determined for all samples in each group at each time point. These concentrations were normalized to the 6 hr value, and changes plotted as a function of time. The mean and SEM normalized change of all metabolites was calculated for each group at each time.

Figure 3 demonstrates metabolites with significant differences (p <0.05, after multiple test corrections) among all the test groups as determined by ANOVA at 6 hr (Figure 3A), and *t*-test analysis of the 7B2-PNU versus no treatment group at 24 hr (Figure 3B). At 6 hr, the levels of 16 intracellular metabolites were significantly affected, and in all but 3 (aspartate, lactate and UMP), metabolite concentrations in the 7B2-dgA treated cells were distinct from the other two groups. The most notable difference included elevated levels of 9 intracellular amino acids in 7B2-dgA treated cells. Of particular note are the branched chain amino acids (BCAAs) leucine, isoleucine and valine, which are known to be elevated during activation and differentiation of immunocytes and T cells in particular (33–37). At six hr all three were significantly elevated in the 7B2-dgA treated cells compared to untreated; and two of the three (leucine and valine) were significantly elevated in the 7B2-PNU treated cells. Levels of creatine, taurine, and fumarate were lower in 7B2-dgA treated cells compared to the 7B2-PNU and no treatment groups. The observation that a significant portion of 7B2-dgA induced changes led to increased levels of intracellular metabolites is evidence that membrane leakage was not the cause of changes observed at this early time point. Both 7B2-dgA and 7B2-PNU treatments resulted in increased intracellular lactate concentrations. Comparison of 7B2-PNU treated and untreated cells at 24 hr (Figure 3B) identified 14 metabolites whose concentrations were significantly different. Many of the metabolite changes mirrored those observed at 6 hr following 7B2-dgA exposure, including increased levels of amino acids, and decreased taurine. In contrast maleate, which was increased 6 hr post 7B2-dgA, was decreased 24 hr post 7B2-PNU. Both 7B2-dgA treated cells at 6 hr and 7B2-PNU treated at 24 hr were in the early stages of cell death, as observed by morphologic examination and trypan blue staining. It is possible that the changes in metabolite concentrations observed at those time points result in part from cell death processes.

**Fig. 3.**
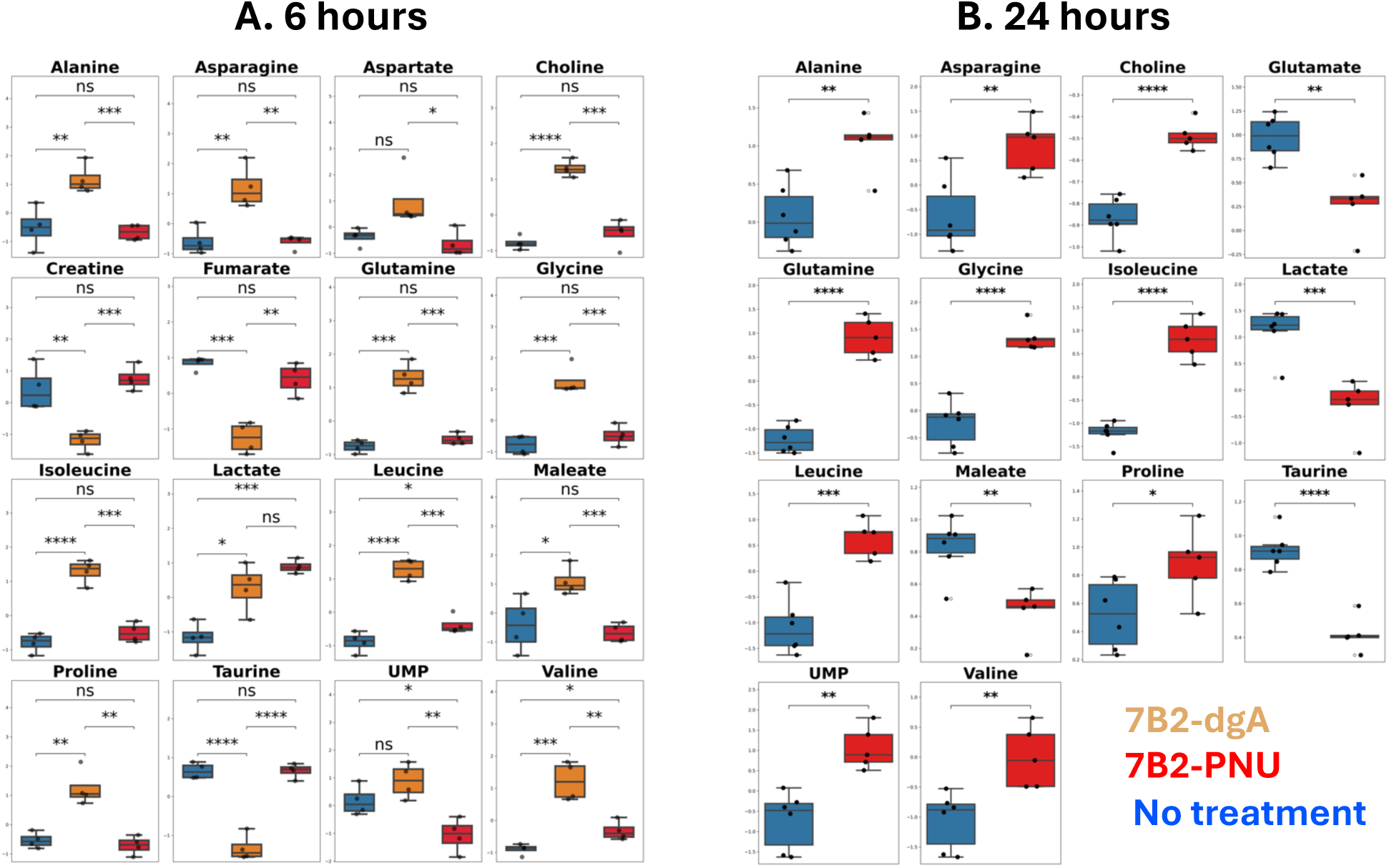
Intracellular metabolites with significantly altered concentrations across treatments. Analyses were performed with an in-house Python workflow (pandas, statsmodels, SciPy, statannotations, matplotlib). Data was normalized to mean, log transformed and autoscaled. **A. Analysis of 6 hr samples.** Metabolites shown met significance by one-way ANOVA across groups (No treatment, 7B2-dgA, 7B2-PNU) with Benjamini–Hochberg FDR correction (q < 0.05). **B. Analysis of 24 hr samples.** Pairwise comparisons between groups are annotated using two-sample t-tests (two-sided. Boxplots show median and IQR with jittered points for individual samples), ns = not significant; * p < .05; ** p < .01; *** p < .001; **** p < .0001.

### Overlapping and distinct transcriptional pathways were activated following treatment of infected cells with ADC and IT

RNA-Seq was used for transcriptional analyses. Total RNA was extracted from cells 6 and 24 hr post treatment with CIC. At 6 hr there was little cytopathic effect (measured as trypan blue exclusion) in any culture, but by 24 hr viability in the 7B2-dgA treated culture was <10% (as measured by trypan blue exclusion). Even in the cultures with few viable cells, RNA yields were robust and of high quality (Supplementary Figure S1A). cDNA libraries were derived from the extracted RNA and sequenced, resulting in a mean of 4.1 X10^7^ reads per sample, >91% of which were >30 bp, a DNA quality score of 35.85 (between10^-3^ to 10^-4^ errors/bp) for a yield of 14,000 Mbp per sample. Greater than 90% of unique reads mapped to the reference human genome GRCh38. “Hit” lists of genes mapped to the human transcriptome were created and normalized to transcripts per million (TPM, Supplementary Table S2). We restricted our analyses to transcripts present in quantities >10 TPM and a variance <15%, which allowed the analysis of 8616 unique transcripts.

Our initial analyses of the data revealed the PCA shown in Supplementary Figure S1B. Each group showed a single widely aberrant point. The aberrant points were found in samples from the wells in the column on the right border of each plate (for both 6 and 24 hr samples). Visual inspection (Figure S1C), cell counts, RNA yield or sequence quality analyses failed to pinpoint a difference between these wells and their replicates. Because the error was clearly systematic, even though we could not identify the reason, we eliminated these samples from the analyses shown here. PCA analysis of transcript profiles (Figure 4A) demonstrated that at 6 hr the 7B2-dgA treated cells diverged from 7B2-PNU and untreated cells, and that by 24 hr each treatment group clustered separately from the others based on distinct transcriptomic profiles. This temporal pattern was similar to that observed for the metabolite changes (Figure 2). At 24 hr, cells treated with 7B2-dgA were <10% viable, thus it is likely that these data represent the late transcripts of dying cells. We next submitted samples at each time point to two-way analyses using DESeq2 to identify genes with significantly different expression levels between the two groups. Significance is defined as >2-fold change plus padj<0.05. Figure 4B summarizes the results. Supplementary Table S3 presents the complete listing of all the significant genes for each comparison, their fold change and p value adjusted for the 8616 comparisons made (padj). Consistent with the PCA, no differences were observed between untreated and 7B2-PNU treated cells at 6 hr, whereas 7B2-dgA treated cells differed from the other two groups. At 6 hr the majority of differentially expressed genes were up-regulated in the 7B2-dgA treated group compared to the others, as demonstrated by the volcano plot shown in Figure 4C, which represents a comparison of untreated vs 7B2-dgA treated cells at this time point. Many of the most upregulated genes were transcription factors associated with cell death and/or growth arrest pathways, including EGR1, 2, and 4, BTG2, DUSP2, JUN, JUNB, and NR4A1 and 2 (38–44). Expression of these genes was also elevated at 24 hr in both treatment groups, compared to untreated cells. At 24 hr, a much larger set of genes was differentially regulated, approaching 25% of the total transcripts when comparing the untreated versus 7B2-dgA treated cells. Additionally, compared to untreated cells, both treatments resulted in more upregulated than downregulated transcripts at 24 hr, although not as striking as in the 7B2-dgA treated group at 6 hr. Among the genes upregulated at 24 hr are members of the growth arrest specific gene family, GAS5 and GAS8.

**Figure 4.**
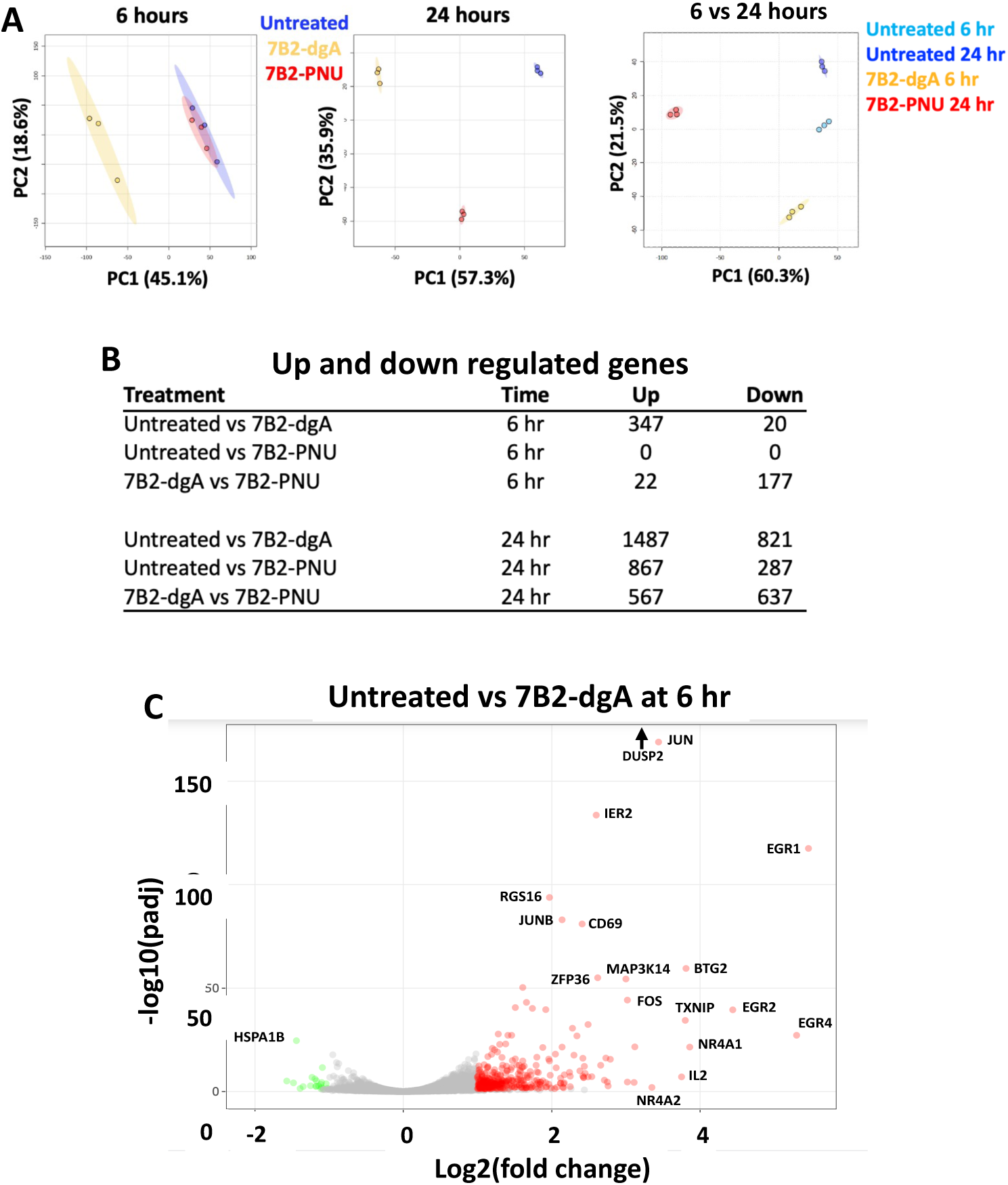
Analysis of individual transcripts by RNA-Seq. A. 2D-PCA scores plot. Transcriptional data were uploaded to ExpressAnalyst, the PCA analysis was performed on data normalized by TPM and log transformed using the DESeq2 module within it. **B. Number of significantly altered transcripts.** Pairwise comparisons of treatment groups performed with DESeq2. Gene expression was considered significantly altered if padj value was <0.05 and absolute log_2_ fold change >1. **C. Volcano plot** of significantly altered transcripts in 7B2-dgA treated cells compared to untreated cells at 6 hr. Upregulated are in red, downregulated in green.

We have noted that killing was more rapid with 7B2-dgA than 7B2-PNU, Figure 1, and references (1, 17). Therefore we explored whether the differences in gene expression between the two CICs at each time point were simply a reflection of these kinetic differences. To test this possibility, we performed a PCA, comparing gene expression 6 hr post 7B2-dgA and 24 hr post 7B2-PNU treatments (Figure 4A, right). The analysis was complicated by changes in the untreated samples at the two time points, likely reflecting differences in cell growth over time as the tissue cultures matured. However, despite the changes in the untreated samples, there remained distinct differences between the 6 hr 7B2-dgA and 24 hr 7B2-PNU treated cells, manifest by group separation along both PC1 and PC2 (together accounting for 81.8% of the variance), and indicating that the difference between 7B2-dgA and 7B2-PNU was not only a matter of kinetics, but also included differential gene expression. We quantified the number of HIV transcripts at 24 hr post treatment (Table I), but found no consistent differences among the groups, suggesting that HIV transcription continued even in dying cells. The specificity of the search for HIV transcripts was demonstrated by the very low level of matches obtained in uninfected H9 cells. The few matches observed likely reflect some degree of sequence homology between endogenous retroviruses and the HIV search sequences used.

**Table 1.** HIV Transcripts Identified by Homology Search.

| **Rx<sup>b</sup>** | **Gag<sup>a</sup>** |  | **Env** |  | **5' Nef** |  | **3' Nef** |  |
| --- | --- | --- | --- | --- | --- | --- | --- | --- |
|  | **Mean** | **SEM** | **Mean** | **SEM** | **Mean** | **SEM** | **Mean** | **SEM** |
| **None** | 1274<sup>c</sup> | 105 | 843 | 86 | 1650 | 168 | 1174 | 145 |
| **7B2-dgA** | 1766 | 250 | 586 | 14 | 1998 | 295 | 1449 | 220 |
| **7B2-PNU** | 1147 | 77 | 551 | 28 | 1470 | 117 | 1077 | 90 |
| **H9 cells** | 49 | 2 | 40 | 5 | 98 | 25 | 50 | 4 |

<sup>a</sup> Sequences used to identify transcripts (numbering based on H9/NL4-3 variant provirus sequence, accession AF070521): Gag: 641-740, Env: 6401-6500, 5' Nef: 8777-8876, 3'Nef: 9281-9379.
<sup>b</sup> Sequences obtained from 24 hr post-treatment samples
<sup>c</sup> HIV transcripts per million

We next performed gene pathway analysis, identifying differentially expressed pathways by the number of genes significantly altered relative to the full complement of genes in that pathway. There exist multiple classification schemes of gene pathways; here we discuss KEGG pathway analyses, but Supplementary Tables S4-S7 contain the results from KEGG, Reactome, GO Process, GO Component, and GO Function identification algorithms. Table 2 reports up and downregulated KEGG pathways, listed by false discovery rate, most significant at top, eliminating cell-type irrelevant pathways, and truncating the listing at 20, if necessary. Supplementary Tables S4-S7 provide the full list of significantly altered pathways with a list of the genes in that pathway whose expression was significantly altered, as well as padj of the false discovery rate. In comparison to the untreated control, by 6 hr 7B2-dgA induced transcription of genes within the apoptosis pathway, and upregulated gene transcripts from several important signaling pathways including TNF, MAPK, p53 and NF-κB. The expression of these pathways remained upregulated at 24 hr. The single pathway downregulated 6 hr following 7B2-dgA treatment was protein processing in the endoplasmic reticulum (ER), perhaps in response to ribosomal inactivation by ricin. 24 hr after 7B2-dgA exposure, a number of metabolic pathways were downregulated, although ribosome biogenesis was activated, even during this terminal state. Apoptosis pathways were upregulated at both 6 and 24 hr following 7B2-dgA, while ferroptosis was downregulated at 24 hr. By 24 hr, 7B2-PNU treated cells had altered many of the pathways observed 6 hr following 7B2-dgA treatment indicating a common response to a death signal: apoptosis and upregulation of TNF, NF-κB, p53 signaling. Other similarities included the late downregulation of metabolic pathways. But there were also differences in the responses 24 hr following 7B2-dgA and 7B2-PNU, highlighted in the rightmost column of Table 2. DNA replication pathways were upregulated following 7B2-PNU, but not 7B2-dgA. It is an apparent contradiction of these analyses that some pathways appear to be both upregulated and downregulated simultaneously, if a sufficient number of up and down regulated transcripts in the pathway were found to be significantly altered (for example, cell senescence in the 24 hr comparison between control and 7B2-PNU). While this may represent a statistical quirk, it could also reflect a true perturbation of the pathway. Figure 5 presents heat maps of specific transcripts within key pathways following 24 hr of treatment. Both similarities and differences in pathway activation were observed among the treatment groups. Consistent with the PCA (Figure 4A), each treatment group mapped distinctly from the others at 24 hr. The similarities between 7B2-dgA and 7B2-PNU treated groups were much greater than that between either CIC and untreated cells. Even among the pathways activated following treatment with either CIC (apoptosis and signaling by TNF, NF-κB, and p53), there remained differences in activation among sub-groups of genes though there was great overall similarity. Other pathways appeared distinctly different following treatment with each CIC: ubiquitin mediated proteolysis, spliceosome, and pathways in cancer.

**Figure 5.**
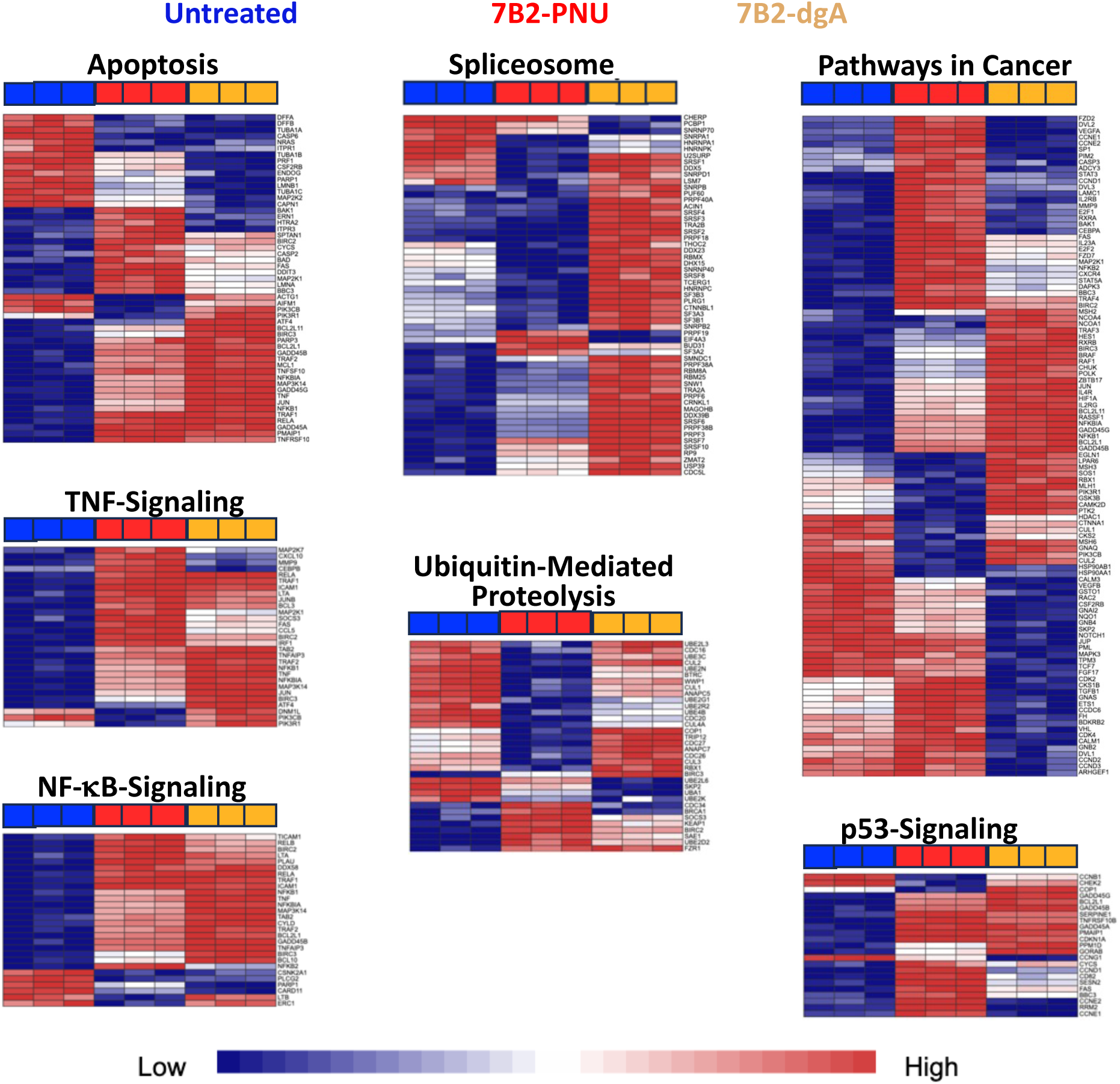
Heat maps of selected gene pathways significantly altered following CIC treatment. ExpressAnalyst software used GSEA to identify KEGG gene pathways significantly altered 24 hr following treatment with CIC and created heat map representations of gene expression. The results of individual triplicates in each treatment group are shown. Heat maps were organized using Ward’s method and ranked vertically by padj (most significant at the top). Treatment groups are indicated above each map. As demonstrated, the samples in each treatment group map together. Specific genes are listed on the right and names can be viewed under high zoom. The expression level for each gene in each sample is shown in the horizontal rows. Expression levels are shown as ranging from low (blue) to high (red).

**Table 2.**
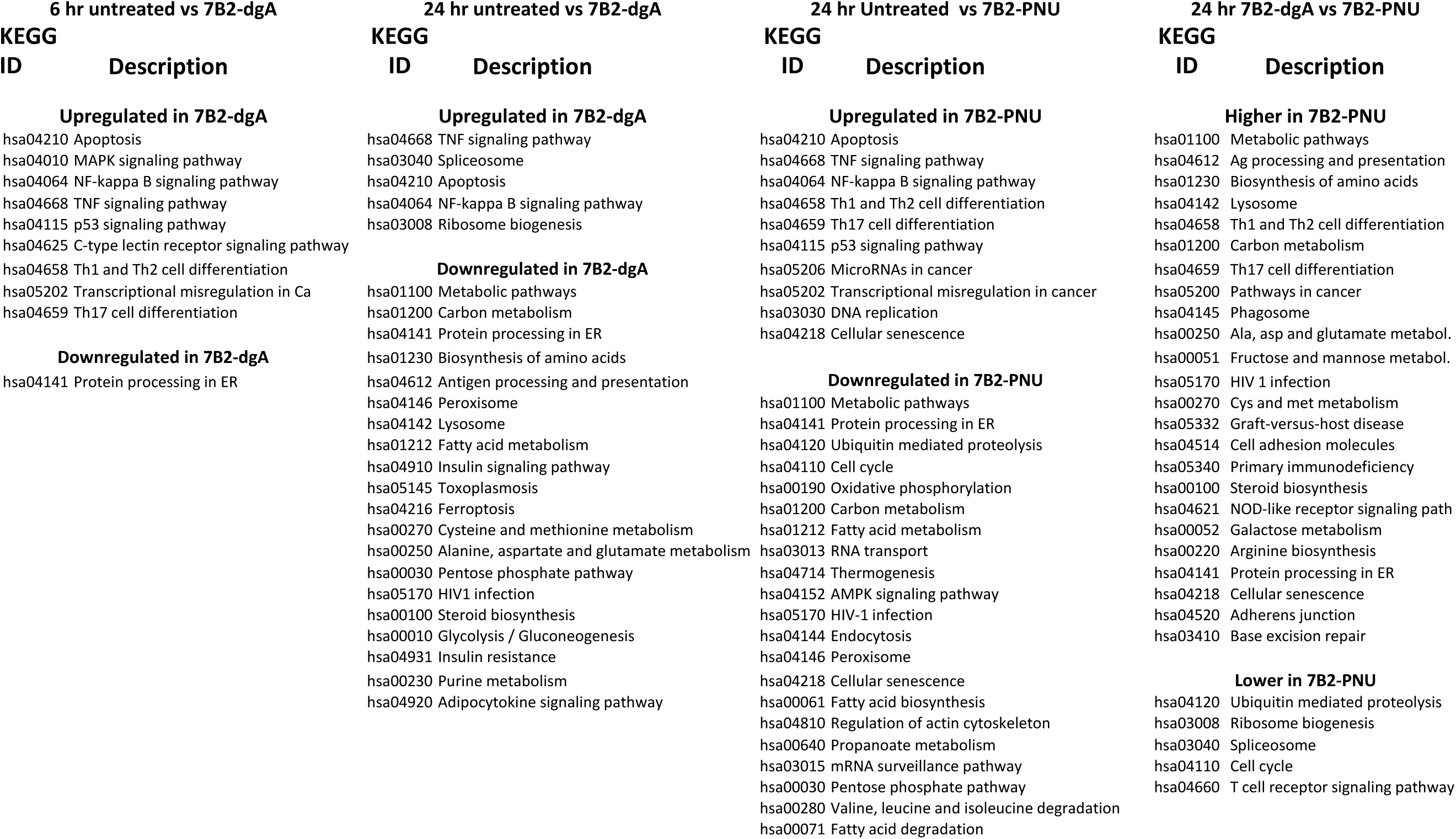
Differentially Expressed KEGG Pathways.

## DISCUSSION

We have investigated the effects of two different CICs, an IT (7B2-dgA) and an ADC (7B2-PNU), each targeted by the same anti-gp41 mAb (7B2) to kill HIV-infected CD4+ lymphoma H9/NL4-3 cells. We evaluated the composition of intracellular metabolites and the transcriptome. These parameters are influenced by current conditions within the cell, change dynamically as cells undergo alterations, and are integrally related to one another, i.e. the metabolic status is influenced by the enzymatic pathways under transcriptional regulation. Because cell death is associated with the release of various proteases, proteomic analyses were deemed less likely to provide interpretable information.

The CICs each carried 1-3 toxic molecules per antibody, but differed in their linkers (1). In a companion paper, we showed that 7B2-dgA induced growth arrest and cytotoxicity more rapidly than 7B2-PNU, was more effective at lower concentrations, and required shorter exposure to produce effects. 7B2-PNU killed Env-negative bystander cells, and surprisingly caused growth enhancement upon low dose or short-term exposure (17). The analyses reported here showed many shared effects of CIC treatment on the cells, reflecting in part their status as dying cells. But the CICs also had unique effects, likely reflecting their mode of cellular injury. Understanding the mechanisms whereby different CICs kill their targets may have clinical relevance in determining the specific cytotoxic payload for a particular use. Although these studies directly address the use of CICs to eliminate HIV-infected cells, they are also relevant for the design of CICs in treating cancer, other persistent viral infections, autoimmunity and other illnesses.

The effects of ricin and ricin-containing ITs on cells are well studied (45–48). Ricin A chain functions as an N-glycosidase, inactivating the large ribosomal subunit, resulting in a cessation of protein synthesis, and inducing apoptosis. Ricin or IT is internalized into endocytic vesicles, where the free A chain is released by reduction. The A chain is retro-transported via the Golgi and ER, into its site of action, the cytoplasm. There one A chain can cleave 3000 ribosomes per minute. Our observations of increased intracellular pools of some amino acids, downregulated protein processing pathways in the ER early after treatment with 7B2-dgA, and increased ribosome biogenesis later are all consistent with ricin’s main mechanism of action, inhibition of protein synthesis, and attempts to counter it. Induction of apoptosis is well described following ricin exposure, as are activation of MAPK, p53, NF-κB, and TNF signaling pathways. We found little evidence that other death pathways were activated, as has been reported for ricin (48–50). 7B2-dgA’s rapidity of action, potency and ability to deliver a lethal hit in a brief time, most likely reflects the potent enzymatic activity of ricin A chain.

PNU-159682 is an anthracycline chemotherapeutic, a metabolite of nemorubicin, produced by microsomal CYP3A-mediated oxidation of nemorubicin in the liver (51). PNU is >6000X more cytotoxic than doxorubicin (1, 52). ADCs constructed with PNU have been tested in preclinical and early clinical trials (52–57). Anthracyclines function by intercalating into DNA and inhibiting topoisomerase II (58), and PNU most likely exerts its cytotoxic effect in this manner as well (53). But given its extreme toxicity, it has been suggested that PNU also affects other cellular functions, in addition to topoisomerase II inhibition (59). The upregulation of KEGG pathways that are unique to 7B2-PNU (ie not elevated following 7B2-dgA), for example, DNA replication, transcriptional misregulation and cellular senescence may offer clues to alternate pathways of cellular injury by PNU. The transcriptional upregulation of the DNA replication pathway following 7B2-PNU treatment may explain our earlier observation that sublethal doses of 7B2-PNU enhanced cellular replication (17). Because PNU was bound to the antibody by a non-cleavable linker, protease digestion of the antibody was required before free PNU is released (52). This extra step may, in part, account for the delayed cytotoxic effect of 7B2-PNU compared to 7B2-dgA.

The metabolic and gene expression studies were performed on persistently-infected H9/NL4-3 cells progressing through different stages of cell death. We observed many commonalities among the alterations following treatment with 7B2-dgA or 7B2-PNU, although with different kinetics. Six hr following 7B2-dgA treatment, cell growth slowed and cells displayed signs of distress, such as altered size, shape, and fragmentation, and by 24 hr, almost all cells were dead and/or dying. Surprisingly the RNA isolated from cells 24 hr post 7B2-dgA was of high quality and yielded excellent sequencing results. This RNA likely represents the last transcripts of dying cells. Similar cell growth, viability, and morphology was observed following 7B2-PNU treatment, but at later time points.

Transcriptional changes were observed 6 hr following 7B2-dgA treatment. These included activation of apoptosis, TNF-signaling, as well as canonical cell-activation and differentiation pathways (Table 2). Similar changes were observed following 7B2-PNU treatment, but only after 24 hr (Table II, Supplementary Tables S3-S7). In the metabolite profiling analyses (Figures 2 and 3), elevated levels of many intracellular AAs were noted 6 hr following 7B2-dgA exposure. Similar but less extensive increases in intracellular AA levels were not observed until 24 hr of 7B2-PNU treatment.

Of the potential cell-death pathways evaluated by RNA-Seq, only apoptosis appeared to play a prominent role following treatment with both the IT and the ADC (Table 2, Supplementary Tables S3-S7). The single pathway downregulated 6 hr following 7B2-dgA treatment was protein processing in the ER, perhaps resulting from dgA’s primary action of ribosomal inactivation (46). Twenty-four hr after 7B2-dgA administration, multiple metabolic networks were downregulated, likely a reflection of the cells’ dying state and the cells’ efforts to counter the damage. Many genes with the greatest degrees of upregulation following administration of either CIC were transcription factors, including EGR1, 2, and 4, BTG2, DUSP2, JUN, JUNB, NR4A1 and NR4A2. These factors have broad and pleiotropic effects on a variety of different cellular processes, and have been implicated in apoptosis, cellular senescence, tumor suppression and other aspects of carcinogenesis (38–44). This pronounced activation of transcription factors resulted in the large numbers of genes whose expression was significantly altered within 24 hr of CIC treatment. The similarities in gene expression between 7B2-dgA and 7B2-PNU treated cells were much greater than that between either CIC and untreated cells, indicating transcriptional commonalities underlying cell death.

A coincident increase in intracellular branched chain amino acids (BCAAs) and the expression of canonical gene pathways associated with T-cell activation and differentiation was observed following both treatments. This suggests that the responses to dying and immune activation share similarities in this T-cell lymphoma line. The accumulation of intracellular BCAAs has been observed during the activation and/or differentiation of T cells and other immunocytes and has been reported to elicit additional metabolic alterations (33–37). These include increased cellular uptake of glucose, altered mitochondrial function, and breakdown of BCAAs by branched chain amino transferases. We observed none of these reported BCAA-induced alterations in our metabolite or RNA-Seq studies, indicating a clear point of divergence among pathways leading to either activation or death.

There were also observable differences in the responses to the two different CICs, beyond those due to kinetic differences. This is best demonstrated by the PCA analyses and heatmaps shown in Figures 4A and 5. The PCAs revealed no overlap of 7B2-dgA and 7B2-PNU treated groups at either time point, or across the two time points. The heatmaps confirmed the PCA results in that each treatment group clustered separately, and further demonstrated patterns where each treatment resulted in similar alterations in gene expression, but also in the expression of genes that were characteristically different for each CIC.

An important goal of new anti-HIV therapies is the long-term suppression of infection allowing patients to remain free from anti-retroviral therapy (ART) for extended periods, i.e. obtain durable remissions. The reservoir of cells carrying a provirus capable of replication, that persists despite years of suppressive ART, is what prevents this outcome. One approach to eliminating these cells is to disrupt latency by activating competent viruses while protecting uninfected cells with ART, sometimes called “activate-and-purge” or “kick-and-kill” (60–63). Evidence of viral activation was observed with different latency disrupting agents, but neither the viral cytopathic effect nor the host immune response caused a decline in the reservoir that was clinically meaningful. More recently, latency disruption in combination with broadly neutralizing mAbs have shown some effects on the reservoir, with mAbs likely functioning through Fc-mediated cytotoxic effects (64–67). We are utilizing CICs to further enhance the ability of anti-HIV mAbs to kill HIV-infected cells. While the downregulation of the KEGG pathway labeled “HIV infection” by both 7B2-dgA and 7B2-PNU at 24 hr (Table 2) may suggest that the CICs have anti-HIV effects beyond killing infected cells, this is balanced by the observation that at 24 hr HIV transcripts were found in equal numbers in cells treated with either or no CIC (Table 1).

In our comparisons of 7B2-dgA and 7B2-PNU, the IT has shown unique advantages over the ADC. But until the problem of immunogenicity can be mitigated, the ADCs have the edge. Considerable effort has gone into solving the problem of IT immunogenicity, with some success. Approaches to minimizing immunogenicity include genetic modification of the IT (21, 22), PEGylation (5, 18), and both specific and nonspecific immunosuppression (19, 20, 24). When the goal of de-immunizing ITs is achieved, our data suggest that ITs may have unique advantages and that their utility may need to be reconsidered.

## ACKNOWLEDGEMENTS

We thank Dr. Jennifer Thomson for showing us the methods for RNA purification and quantification. Dr. Ellen Vitetta read the manuscript and provided meaningful comments. This work was funded by NIH grant R01AI136758, and by the Montana State University NMR Core: NIH SIG program (Grant No. 1S10RR13878 and 1S10RR026659), the National Science Foundation (Grant No. NSF-MRI:DBI-1532078, NSF-MRI:CHE-2018388), the Murdock Charitable Trust Foundation (Grant No. 2015066:MNL), and MSU’s Vice President for Research and Economic Development.

## DATA AVAILABILITY

With the exception of RNA-Seq, all data are included with the manuscript and Supplementary Data. The RNA-Seq raw and processed data are available at the GEO database (Gene Expression Omnibus, National Center for Biotechnology Information), entry number: GSE289396.

## List of Supplementary Material (provided as separate files)

**Supplementary Figure S1.** Preliminary analysis of RNA-Seq data.

**Supplementary Table S1.** Concentrations of intracellular metabolites

**Supplementary Table S2.** RNA-Seq Hit Lists

**Supplementary Table S3.** Differentially Expressed Genes

**Supplementary Table S4.** Differentially expressed gene pathways untreated vs 7B2-dgA 6 hr

**Supplementary Table S5.** Differentially expressed gene pathways untreated vs 7B2-dgA 24 hr

**Supplementary Table S6.** Differentially expressed gene pathways untreated vs 7B2-PNU 24 hr

**Supplementary Table S7.** Differentially expressed gene pathways 7B2-dgA vs 7B2-PNU 24 hr

