## Supplementary Figure S1 for "Anti-HIV Immunotoxin and Antibody-Drug Conjugate Display Both Common and Distinct Effects in Killing Target Cells"

**A**

| Time | CIC | Total RNA (μg) | | RNA Integrity (RIN) | | Total reads | | Unique Mapped Reads | | Percent |
|---|---|---|---|---|---|---|---|---|---|---|
| | | Mean | SEM | Mean | SEM | Mean | SEM | Mean | SEM | Mapped Reads |
| 6 hr | None | 17.37 | 1.69 | 9.63 | 0.03 | 4.31E+07 | 5.74E+06 | 3.98E+07 | 5.23E+06 | 92.40 |
| 6 hr | 7B2-dgA | 11.38 | 2.01 | 9.6 | 0.06 | 4.77E+07 | 5.49E+06 | 4.42E+07 | 4.92E+06 | 92.64 |
| 6 hr | 7B2-PNU | 12.43 | 1.4 | 9.65 | 0.04 | 3.48E+07 | 3.10E+06 | 3.22E+07 | 2.98E+06 | 92.49 |
| 24 hr | None | 20.13 | 0.93 | 9.65 | 0.05 | 4.01E+07 | 4.10E+06 | 3.66E+07 | 3.84E+06 | 91.27 |
| 24 hr | 7B2-dgA | 7.4 | 2.86 | 8.7 | 0.31 | 3.63E+07 | 2.84E+06 | 3.33E+07 | 2.78E+06 | 91.79 |
| 24 hr | 7B2-PNU | 13.53 | 1.24 | 9.1 | 0 | 3.61E+07 | 4.28E+06 | 3.32E+07 | 3.95E+06 | 92.01 |

**B**

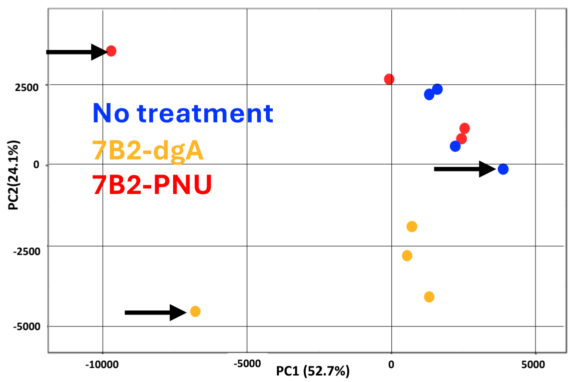


**C**

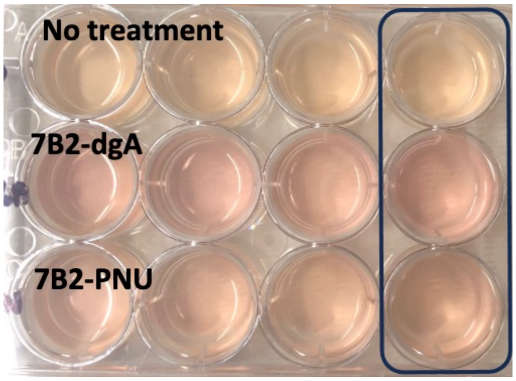


**Supplementary Figure S1. Preliminary analysis of RNA-Seq data. Panel A.** RNA yield/integrity and library quality. **Panel B.** PCA analysis including all four individual samples from each treatment group at the 6 hr time point. Arrows indicate the samples obtained from the rightmost column of the tissue culture plate. **Panel C** shows the tissue culture plate immediately prior to harvest, with the aberrant cultures shown within the box.
